# Inflammation mediates lung cancer-induced increase in the epidermal green autofluorescence of mice

**DOI:** 10.1101/2020.09.02.259200

**Authors:** Mingchao Zhang, Wanzhi Tang, Weihai Ying

## Abstract

Skin’s autofluorescence (AF) holds great promise for non-invasive diagnosis of diseases. Our previous study has indicated that keratin-based skin’s AF holds great promise to become a novel biomarker for diagnosis of multiple major diseases, including lung cancer, acute ischemic stroke, Parkinson’s diseases, stable coronary artery disease and myocardial infarction. It is critical to investigate the mechanisms underlying the increased skin’s AF of the patients. Our previous study has shown that development of lung cancer can induce increased skin’s green AF of mice, while the mechanisms underlying the increased AF have remained unclear. We hypothesized that the inflammation in the mice with lung cancer is causative to the increased AF. Our study found that the anti-inflammation drug indomethacin virtually abolished the lung cancer-induced increase in the skin’s green AF. We further found that the AF was originated from the epidermis of the mice. Our findings have provided critical information for understanding the mechanisms underlying the lung cancer-induced increase in the epidermal green AF, indicating that at least for lung cancer, the disease-induced increase in the skin’s AF mainly results from increased inflammation in the body.

## Introduction

Inflammation is key common pathological factor of a number of major diseases (1–4). Chronic systemic inflammation can also promote development of multiple diseases including cancer (5–7). However, there has been no biomarker for determining non-invasively the levels of inflammation in human body. Due to the critical importance of inflammation, it is of significance to search for the biomarkers of this type.

Human autofluorescence (AF) of the dermis has shown promise for non-invasive diagnosis of diabetes and diabetes-related pathology (8). Our previous study has also indicated a significant promise of skin’s AF to become a novel biomarker for diagnosis of multiple major diseases, including lung cancer (9), acute ischemic stroke (10), Parkinson’s diseases (11), myocardial ischemia (12) and stable coronary artery disease (12). It is critical to investigate the mechanisms underlying the increased skin’s AF of the patients.

Our previous studies have shown that development of lung cancer can induce increased skin’s green AF of mice (9). However, the mechanisms underlying the increased AF have remained unclear. We hypothesized that the inflammation in the body of the mice with lung cancer is causative to the increased AF due to the following information: First, our previous studies have shown that Lipopolysaccharide (LPS), a widely used inflammation inducer, can dose-dependently induce increased skin’s green AF (13); and second, there are marked increases in inflammation in the body of lung cancer patients (14, 15). Our current study has found that the anti-inflammation drug indomethacin can virtually abolish the lung cancer-induced increase in the skin’s green AF, providing strong support to our hypothesis.

## Materials and Methods

### Materials

All chemicals were purchased from Sigma (St. Louis, MO, USA) except where noted. Male C57BL/6SLACBL/6Slac mice of SPF grade were purchased from SLRC Laboratory (Shanghai, China).

### Development of a mouse model of lung cancer

Animal studies were conducted according to an animal protocol approved by the Animal Protocol Committee, School of Biomedical Engineering, Shanghai Jiao Tong University. Cell suspension of LLC (Lewis Lung Carcinoma) cells (1 × 10^6^ cells) in a total volume of 5 μl mixed with Matrigel (PBS : Matrigel = 4:1) were injected into the left lung of 4-week-old male C57BL/6 mice. The mice were sacrificed nine days after the injection, and their lungs were obtained for determining if lung cancer was developed.

### Imaging of the AF of mouse’s skin

The skin’s AF of the ears of the mice was imaged by a two-photon fluorescence microscope (A1 plus, Nikon Instech Co., Ltd., Tokyo, Japan), with the excitation wavelength of 488 nm and the emission wavelength of 500 - 530 nm. The AF was quantified as described previously (13).

### Statistical analyses

All data are presented as mean + SEM. Data were assessed by one-way ANOVA, followed by Student - Newman - Keuls *post hoc* test, except where noted. *P* values less than 0.05 were considered statistically significant.

## Results

We determined the effects of the anti-inflammation drug indomethacin on the development of lung cancer of the mice. Nine days after injection of LLC, all of the mice developed lung cancer (Fig. 1A). There was no significant difference between the weight of the lung of the mice that were injected with LLC only and the weight of the lung of the mice that were injected with both LLC and indomethacin (Fig. 1B).

**Fig. 1.**
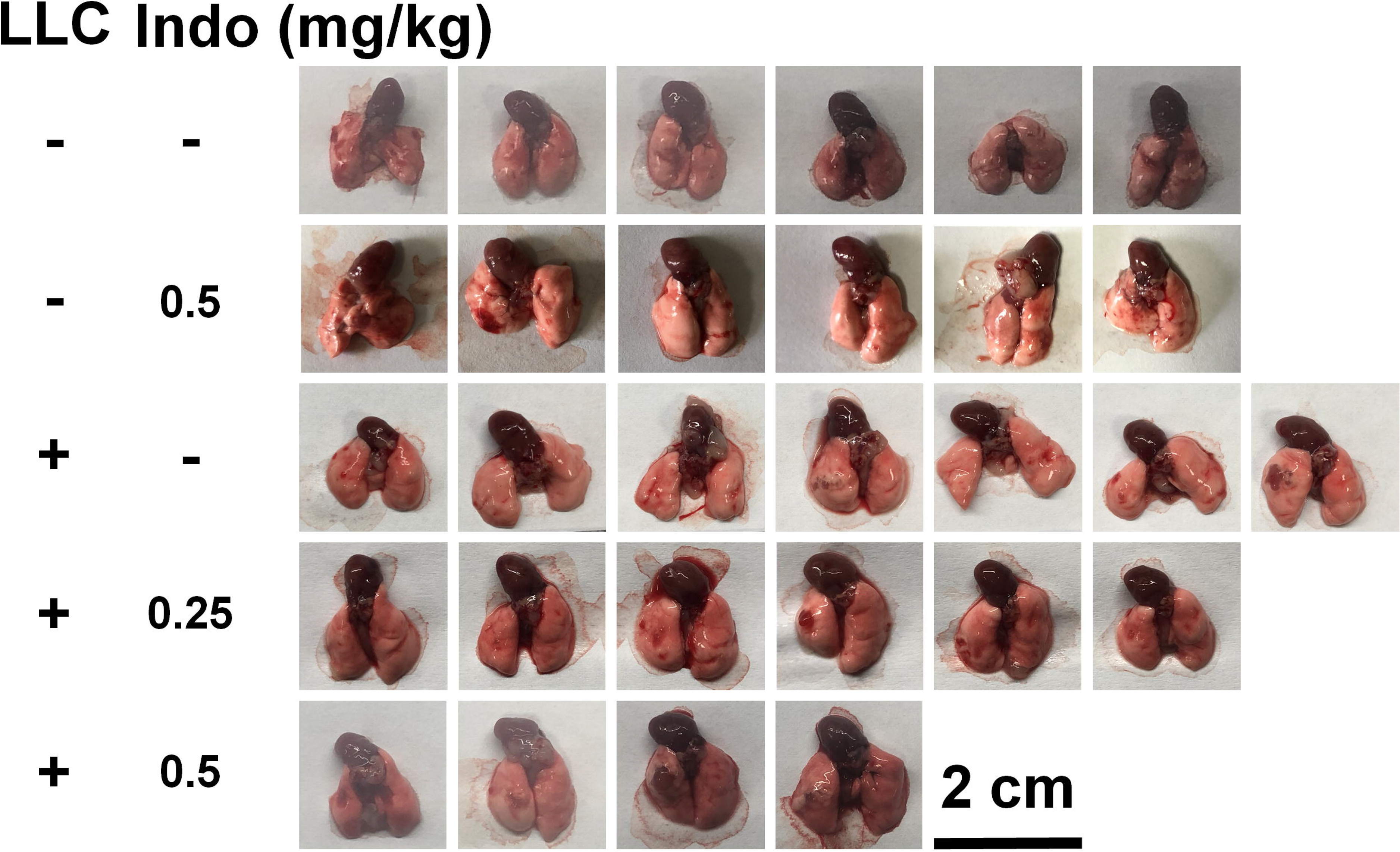
Effects of Indomethacin (Indo) on the weight of the lung in the mouse model of lung cancer. (A) One week after injection of LLC, all of the mice developed lung cancer. (B) There was no significant difference between the weight of the lung of the mice that were injected with LLC only and that of the mice that were injected with both LLC and Indo. Three days after the mice were injected with LLC, Indo at the doses of 0.25 or 0.5 mg/ml was injected. Nine days after the LLC injection, imaging of the skin’s green AF of the mice was conducted. After sacrifice of the mice, the lungs of the mice were imaged by a camera, and the lungs of the mice were also weighted. N = 4 - 7. ##, *P* < 0.01 (Student t-test); *, *P* < 0.05; **, *P* < 0.01.

We further determined the effects indomethacin on the lung cancer-induced increases in the skin’s green AF of the mice: Development of the lung cancer led to a marked increase in the skin’s green AF (Figs. 2A and 2B). Administration of either 0.25 or 0.5 mg/kg indomethacin virtually abolished the lung cancer-induced increase in the AF (Figs. 2A and 2B).

**Fig. 2.**
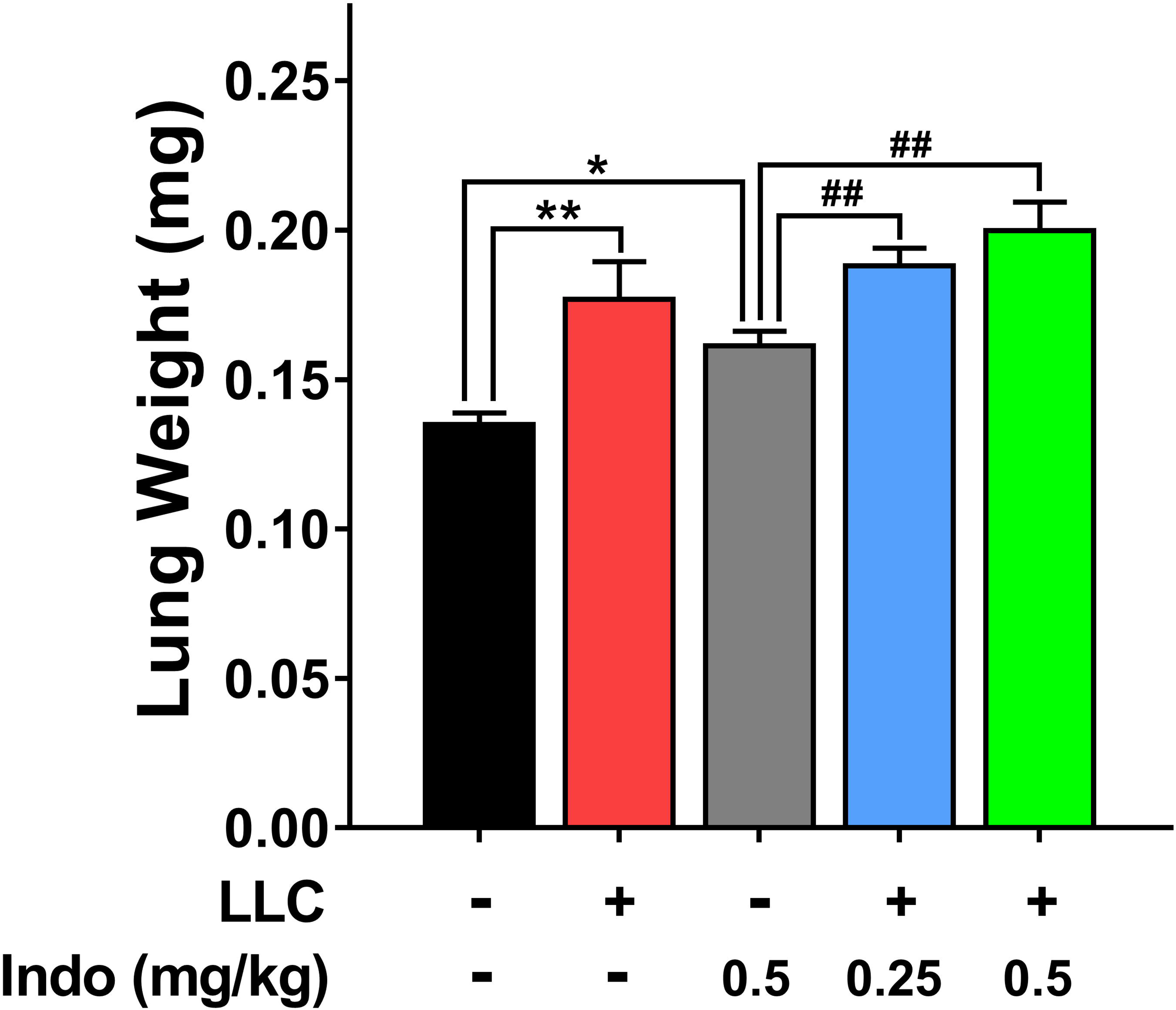
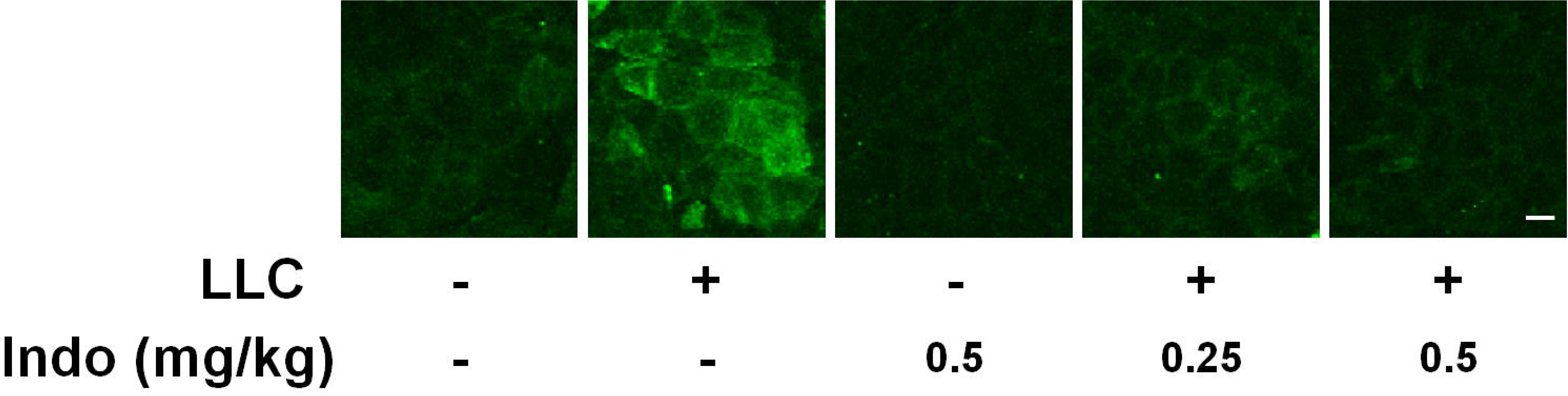
Effects of Indomethacin (Indo) on the skin’s green AF of the mice with lung cancer. (A, B) Development of lung cancer led to a significant increase in the skin’s green AF of the mice. Administration of 0.25 or 0.5 mg/kg Indo virtually abolished the lung cancer-induced increase in the AF. N = 4 - 7. #, *P* < 0.05 (Student t-test); **, *P* < 0.01.

Orthographic green AF images of the mouse’s skin showed that the lung cancer-induced AF occurred only at certain layer of the skin with the thickness of approximately 10 μm, which was approximately 10 - 20 μm apart from the outer surface of the epidermis. This observation indicates that the AF was originated from the epidermis (Fig. 3).

**Fig. 3.**
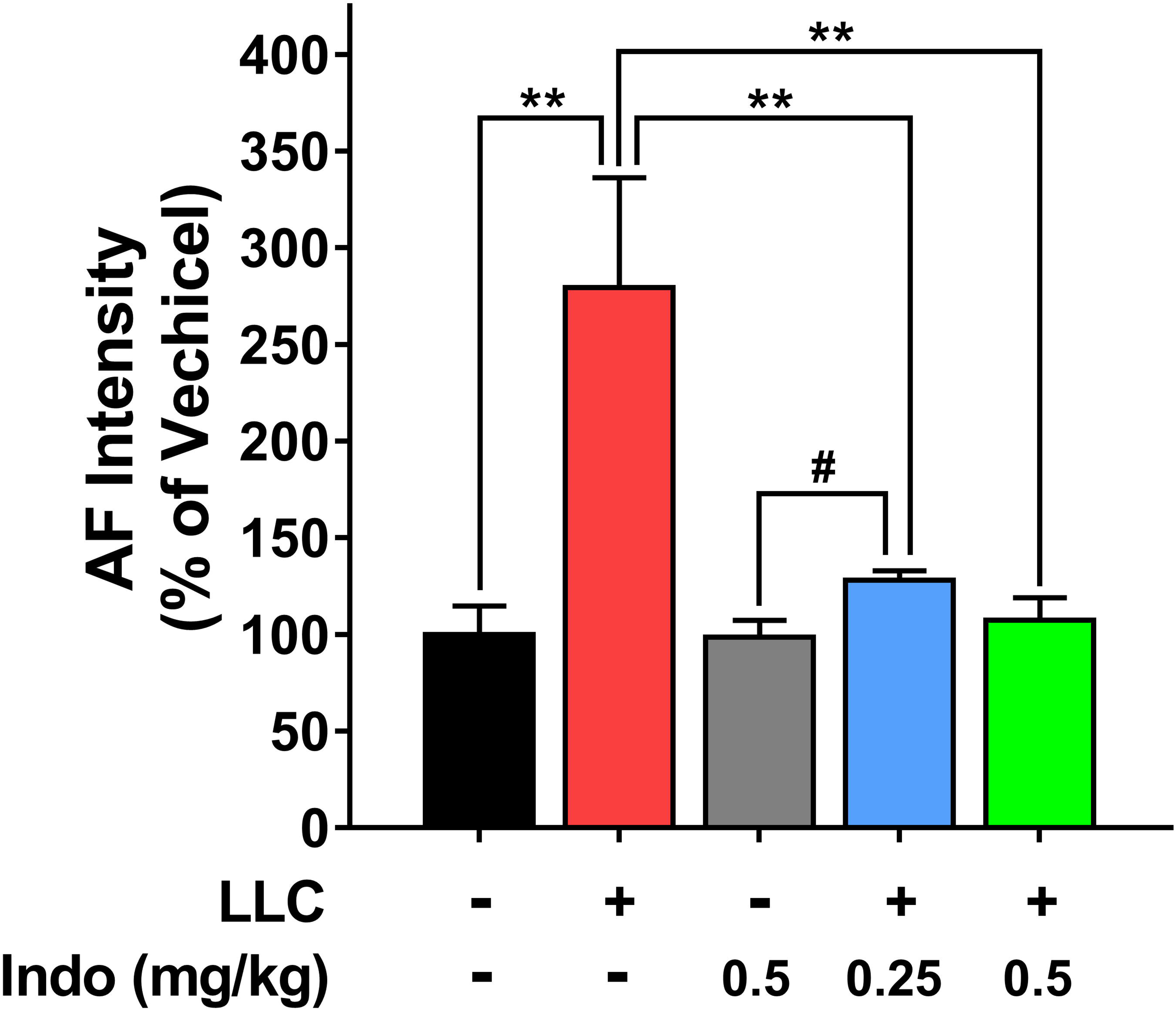
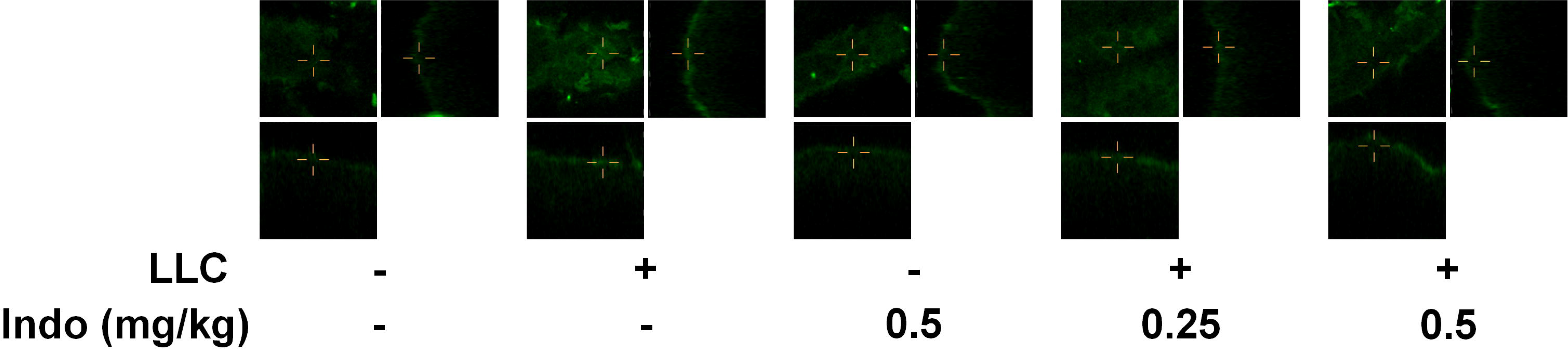
Analyses of the orthographic green AF images indicated that the lung cancer-induced AF was originated from the epidermis of the mice. Orthographic green AF images of the mouse’s skin showed that for both control mice and the mice with lung cancer, the skin’s green AF occurred only at certain layer of the skin with the thickness of approximately 10 μm, which was approximately 10 - 20 μm apart from the outer surface of the epidermis. N = 4 - 7.

## Discussion

The major findings of our current study include: First, the anti-inflammation drug indomethacin virtually abolished the lung cancer-induced increase in the skin’s green AF; and second, the AF was originated from the epidermis of the mice. Our study also showed that indomethacin did not significantly affect the wight of the lung of the mice with lung cancer, suggesting that indomethacin did not significantly affect the size of the lung cancer. This observation has argued against the possibility that the indomethacin-produced blockage of the lung cancer-induced AF increase resulted from its effects on the size of the tumors. Collectively, our current finding has indicated that at least for lung cancer, the disease-induced increase in the epidermal AF mainly results from the increased inflammation in the body.

Our previous study has indicated great promise of skin’s AF to become a novel biomarker for diagnosis of multiple major diseases. It is critical to investigate the mechanisms underlying the increased skin’s AF of the patients. Our previous studies have shown that development of lung cancer can induce increased skin’s green AF of mice (9). Since inflammation can induce increased skin’s AF of mice (13), which has been found in the body of lung cancer patients (14, 15), we hypothesized that inflammation mediates the lung cancer-induced AF increase. Our study has found that indomethacin virtually abolished the lung cancer-induced increased in the epidermal AF, indicating that inflammation mediates the AF increase.

We have found that both inflammation and oxidative stress can induce increased epidermal green AF of mice (13). Since there are both increased inflammation (14, 15) and oxidative stress (16–20) in the body of lung cancer patients, theoretically both inflammation and oxidative stress may contribute to the increased AF. However, indomethacin abolished the lung cancer-induced increased in the AF, suggesting that at least for lung cancer, the disease-induced increase in the AF results mainly from increased inflammation in the body.

Future studies are needed to investigate the mechanisms underlying the disease-induced increase in skin’s AF in other disease models. These studies will be highly valuable for establishing the keratin’s AF-based approaches for disease diagnosis.

## Acknowledgment

The authors would like to acknowledge the financial support by two research grants from a Major Special Program Grant of Shanghai Municipality (Grant # 2017SHZDZX01) (to W.Y.).

